# *biodumpy*: A Comprehensive Biological Data Downloader

**DOI:** 10.1101/2025.07.26.666724

**Authors:** Tommaso Cancellario, Tomas Golomb Durán, Antoni Josep Far, Alejandro Roldán, Maria Capa

**Affiliations:** Centre Balear de Biodiversitat, Universitat de les Illes Balears, Palma, Spain

**Keywords:** biodiversity, data mining, data processing, ecology, genetics, python, taxonomy

## Abstract

In recent years, the expansion of public biodiversity platforms and associated datasets has greatly improved access to ecological and biological information. These resources now cover vast geographic areas, extended temporal scales, and diverse taxonomic groups, becoming essential for ecological studies by enabling more comprehensive analyses and novel hypotheses testing. Concurrently, the development of programming packages has facilitated data access and interaction, streamlining their retrieval processes. However, most existing tools are limited to specific databases, posing challenges for studies requiring seamless integration of data from multiple sources. The growing availability of biodiversity data highlights the urgent need for robust tools to efficiently process, analyse, and interpret ecological and biological information. To address this limitation, we introduce *biodumpy*, a new Python package developed for the retrieval, management, and integration of biological data from various public databases. *biodumpy* provides access to up-to-date and comprehensive datasets spanning genetic, distributional, taxonomic, and bibliographic sources. It includes specialized modules for efficient data retrieval across taxonomic lists, with the capability to process multiple modules simultaneously. By integrating diverse data sources, *biodumpy* enhances data acquisition, providing researchers with a powerful framework for comprehensive analyses and supporting ecological research to tackle complex environmental challenges.

## 1. INTRODUCTION

Ecology serves as a bridge that integrates diverse scientific disciplines and its interdisciplinary nature is essential for understanding the complex interactions between organisms and their environments, as well as the relationships among biological entities (Goring et al., 2014; Odum, 1971). A deep comprehension of the ecological dynamics is crucial not only for advancing ecological theories but also for addressing global pressures such as climate change, habitat degradation, and biodiversity loss (Banks-Leite et al., 2020). Addressing these challenges require the development of robust strategies to catalogue, assess, and predict biodiversity variations along with their associated ecosystem services. A key step toward achieving this goal is the collection and sharing of diverse ecological data that span fine to broad spatial and temporal scales, encompass multiple biological levels (e.g., from populations to communities), and include various data types (e.g., species distributions, genetic information).

In recent years, the scientific community has witnessed a remarkable expansion of public biodiversity platforms and related datasets (Heberling et al., 2021). These digital infrastructures serve as extensive repositories, often containing highly structured biodiversity data encompassing diverse issues, including taxonomy (e.g., Catalogue of Life; Bánki et al., 2024), species distributions (e.g., Global Biodiversity Information Facility; GBIF Secretariat), and genetic information (e.g., Barcode of Life Data System; Ratnasingham et al., 2024). Nowadays, such digital resources are indispensable for many ecological studies, as they enable users to perform comprehensive analyses or test and develop new hypothesis that were once unfeasible due to limited data availability, poor quality, and low variability. Alongside the growth of these datasets, there has also been a notable rise in programming packages, particularly for the R programming language, designed to connect directly with these databases, including a range of functions that simplify access and interaction, simplifying the retrieval process for users. However, these tools are often limited to specific databases, creating challenges for researchers seeking to seamlessly integrate data from multiple sources. The development of a toolbox capable of combining data from various biodiversity databases would significantly enhance data acquisition processes, providing researchers with a robust starting point for comprehensive and interdisciplinary analyses, representing a valuable advancement for ecological research and its capacity to address complex environmental challenges.

Here, we introduce *biodumpy*, a new Python package designed to simplify the process of retrieving diverse biological information from several public databases. With *biodumpy*, users can easily download, manage, and integrate data from different sources, ensuring access to the most up-to-date and comprehensive biological information available. The package includes specialized modules designed to efficiently download biological data for a list of given taxa, with the option to process multiple modules simultaneously. These modules provide ideal access to various types of biological information, including genetic, taxonomic, distributional, and bibliographic databases. In this framework, *biodumpy* will significantly enhance data acquisition by integrating diverse datasets, providing researchers with a robust foundation for comprehensive analyses.

## 2. SOFTWARE TOOL DESCRIPTION

The *biodumpy* package is developed in Python programming language, and its stable version (v.0.1.8) can be installed from the official repository of Python PyPI using the command: `pip install biodumpy`. For those interested in the development version (v.0.1.18) the package is available from TestPyPI with the following command: pip install biodumpy -i https://test.pypi.org/simple/--extra-index-url https://pypi.org/simple/. Interestingly, *biodumpy* can also be used in R through the *reticulate* package (Ushey et al., 2024), which allows seamless integration of Python code within the R environment.

### 2.1 Starting with *biodumpy*

The use of *biodumpy* is intuitive, with a general structure that is consistent across all modules. Below we describe an overview of the main steps to get started with the package:

1. Load the package into your Python environment.
2. Import one or more specific modules needed to retrieve the data.
3. Define a list of taxa (except for the Crossref module, which accepts a list of DOIs).
4. Configure the *biodumpy* module/s with the required parameters and start the download.

Although each module of *biodumpy* is characterized by specific parameters, three parameters are common across all modules: `bulk`, `output_format`, and `sleep`. `bulk` is a Boolean parameter allowing users to customize the data structure according to their needs during the saving process (Fig. 1). When `bulk` is *True*, the information downloaded for each taxon is stored into a single file. This configuration may be useful if the amount of the total data is limited, for consolidating data, and simplifying file management. Whereas, if `bulk` is *False*, the information for each taxon is saved in a separate file each one named as the taxon. This option is useful for detailed analysis, particularly when individual taxon-files are required or when the data for each taxon is extensive.

**Figure 1:**
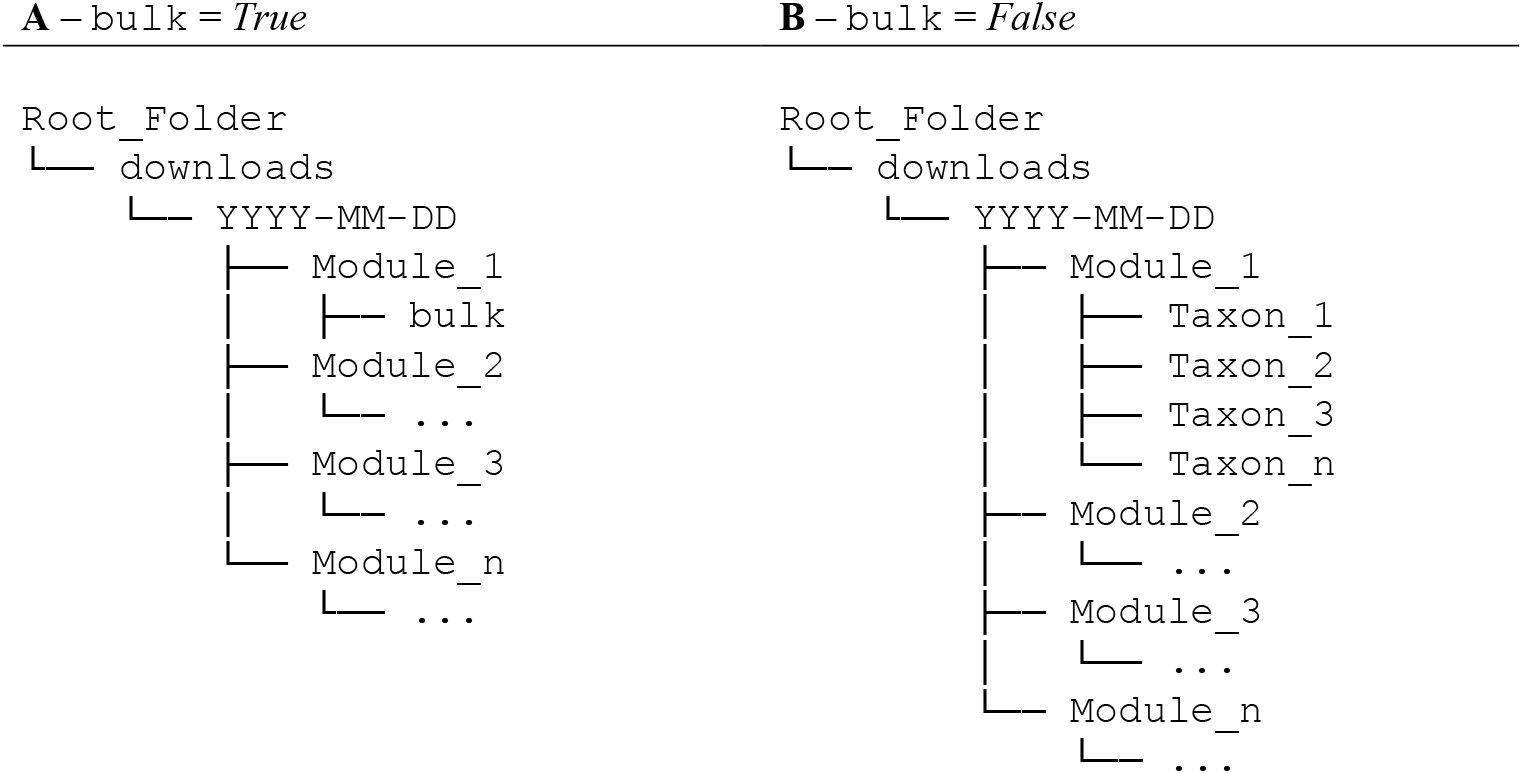
Example illustrating the folder hierarchy structure, with the distinction based on the `bulk` parameter being set to either *True* (A) or *False* (B).

By default, *biodumpy* saves the resulting file in a folder named “downloads” within the user’s working directory. Inside this folder, a subfolder is automatically created, named after the current date and, within it, additional subfolders corresponding to each of the modules used are generated. A general folder hierarchy structure is described in Fig. 1.

Depending on the module, *biodumpy* provides the option to store files in two output formats: JSON or FASTA. Although the JSON format is not commonly used in biological or ecological fields, we have deliberately chosen it as default format due to its flexibility to support nested and complex data structures, such as arrays and key-value pairs. This makes JSON particularly well-suited for representing hierarchical or relational data, such as taxonomy and observations or nested genetic information, within a single file. Additionally, JSON integrates easily with the raw data returned by the Application Programming Interfaces (APIs) used to develop the *biodumpy* modules and can be easily converted into a tabular format (e.g., .csv).

On the other hand, the FASTA format is specifically designed for storing nucleotide or protein sequences. The BOLD and NCBI modules offer the option to download data in FASTA format, making them ideal to retrieve data to perform sequence-based analysis. Finally, the `sleep` parameter is useful for controlling the delay between successive data retrieval requests. It enables users to regulate data download speed, thereby preventing server overloads and avoiding rate limit violations. By default, `sleep` is set to 0.1 seconds. However, we encourage users to adjust this parameter according to the API’s rate policies to ensure responsible data access and avoid overwhelming external servers.

### 2.2 *biodumpy* modules

To date, *biodumpy* includes 10 modules that cater to different aspects of biodiversity data management, providing functionalities for working with a wide range of data types. Below, we describe the main functionalities and parameters for each module. Furthermore, for more detailed information, users can consult the *biodumpy* manual available at https://biodumpy.readthedocs.io/en/latest/.

#### BOLD

The *BOLD* module allows users to retrieve information from the Barcode of Life Data System (Ratnasingham et al., 2024). To select the output format users can simply set the parameter `output_format` to *‘json’* or *‘fasta’*. Additionally, this module provides the option to download a summarized version of the data by setting the `summary` parameter to *True*, offering a more concise and manageable dataset (*Supplementary Material 1 - S1*).

#### COL

The *COL* module allows users to easily retrieve taxonomic nomenclature from the Catalogue of Life (Bánki et al., 2024). This module provides users with the option to verify taxon synonyms by setting the parameter `check_syn` to *True*. When this parameter is enabled, only the accepted name is stored in the classification section of the JSON file. Conversely, both the synonym and the accepted name are included in the classification section (*Supplementary Material 1 - S2*).

#### Crossref

The *Crossref* module allows users to retrieve scientific bibliographic metadata from Crossref database. Unlike other modules that use a list of taxa as an input parameter, this module requires a list of DOIs to download the available bibliographic metadata. Users can also obtain a summarized version of the downloaded data by setting the `summary` parameter to *True* (*Supplementary Material 1 - S3*).

#### GBIF

The *GBIF* module enables users to access taxonomic and distributional data from the GBIF platform (GBIF Secretariat). Taxonomic information can be downloaded using the default parameters, while occurrence information can be obtained by setting the parameter `occ` to *True*. Additionally, the module supports filtering occurrences based on a specified geographic region using the `geometry` parameter. When a geometry is provided, only occurrences that fall within the defined polygon are included in the outcome. Finally, this module allows the user to check for possible nomenclature synonyms using the parameter `accepted_only`. If this parameter is *False*, nomenclature includes possible synonyms; conversely, the information retrieved corresponds only to accepted names (*Supplementary Material 1 - S4*).

#### INaturalist

The *INaturalist* module permit users to retrieve taxon photo metadata from iNaturalist (iNaturalist). Notably, the module does not directly download photos from iNaturalist; instead, it retrieves the metadata required to access them (e.g., 34826202/medium.jpg – photo id and its size). To retrieve a photo, the photo-ID (e.g., 34826202) and the image size (e.g., medium) must be concatenated with the following base URL: https://inaturalist-open-data.s3.amazonaws.com/photos/ (i.e., https://inaturalist-open-data.s3.amazonaws.com/photos/34826202/medium.jpg).

To avoid copyright issues, the module retrieves photos classified under public licenses, including: CC0, CC BY, CC BY-NC, CC BY-NC-ND, CC BY-SA, CC BY-ND, and CC BY-NC-SA (*Supplementary Material 1 - S5*).

#### IUCN

The *IUCN* module enables users to efficiently retrieve information on species assessments, habitats, and threats from the IUCN Red List (IUCN) by specifying a species name and providing a personal API key. Moreover, the module allows users to retrieve data for a specific list of species from one or more areas by specifying a list of regions in the `scope` parameter (e.g., `scope` = *[‘Global’, ‘Europe’]*; *Supplementary Material 1 - S6*).

#### NCBI

The *NCBI* module enables users to efficiently retrieve data from the National Center for Biotechnology Information database (NCBI). To obtain data in FASTA format, users must set the `rettype` and `output_format` parameters to *‘fasta’*.

By providing a list of taxa, users can refine their searches using the `query_type` parameter, which allows for the combination of multiple search criteria to better target the desired data. For example, to download sequences related to the cytochrome c oxidase subunit I, users can set the `query_type` to *‘[Organism] AND (“CO1”[GENE] OR “COI”[GENE] OR “COX1”[GENE] OR “COXI”[GENE])*’ (Porter & Hajibabaei, 2018).

Module performance can be further optimized using the `step_id` and `step_seq` parameters, which control the chunk size for downloading IDs and sequences, respectively. While relatively large chunk sizes are manageable for ID downloads (e.g., `step_id` = *500*), setting a high value for `step_seq` can lead to memory issues due to the potentially large size of the sequence data. To avoid such problems, we recommend using moderate values for the `step_seq` parameter (e.g., `step_seq` = *100*).

Lastly, users can retrieve a summary of metadata associated with the retrieved data by setting the `summary` parameter to *True* (*Supplementary Material 1 - S7*).

#### OBIS

The *OBIS* module facilitates the retrieval of data from the Ocean Biodiversity Information System (OBIS). Users can download taxonomic information from OBIS and, like the GBIF module, this module enables users to download species occurrences by setting the `occ` parameter to *True*. Moreover, the module supports filtering occurrences based on specific geographic regions using the `geometry` or `areaid` parameters. These spatial parameters can be applied independently or in combination. When a `geometry` or `areaid` is specified, the results include only occurrences that fall within the defined polygons (*Supplementary Material 1 - S8*).

#### WORMS

The *WORMS* module enables users to easily retrieve nomenclature information from the World Register of Marine Species database (Ahyong et al., 2024). Users can set the parameter `marine_only` to *True* to restrict the search only for species belonging to the marine environment. Moreover, it is possible to retrieve the distribution data for the taxa setting the parameter `distribution` to *True* (*Supplementary Material 1 - S9*).

#### ZOOBANK

The *ZooBank* module allows users to easily retrieve scientific bibliographic information about taxa from the Official Registry of Zoological Nomenclature database (International Commission on Zoological Nomenclature) by providing a taxa name. To optimize the performance of the module, this function includes the parameter `dataset_size`. This parameter can set to *‘small’* or *‘large’* depending on the amount of data to download. Users can also download additional information setting the parameter `info` to *True*. When this parameter is activated, the outcome includes the main information of a bibliographic resource along with additional details such as the DOI. Examples of retrieved data can be found in *Supplementary Material 1 - S10*.

### 2.3 Customizable models

Although several modules have been implemented in *biodumpy*, some digital sources may remain unexplored, limiting the scope of data retrieval. To address this issue, we offer users the possibility to create their own modules, leveraging the inherent hierarchical structure of *biodumpy*. The commented code to create your own module is available in *Supplementary Material 1 - S11*.

### 2.4 *biodumpy* tests

To further improve the quality and reliability of *biodumpy*, we incorporated a comprehensive suite of tests designed to stress all individual modules and assess their performance under various conditions. We implemented the testing framework using *pytest* (v. 8.3.3) (Krekel et al., 2004). In Table 1, we reported the coverage result for each module.

**Table 1.** Summary the test coverage results for each module. Stmts: Total number of statements (lines of executable code) in the file or module analysed. Miss: Number of statements that were not executed (i.e., not covered by any test). Cover: Percentage of statements that were executed during testing. Missing: Specific line numbers or ranges of lines where statements were not executed.

| Module Name | Stmts | Miss | Cover | Missing |
| --- | --- | --- | --- | --- |
| BOLD | 93 | 0 | 100% |  |
| COL | 69 | 0 | 100% |  |
| Crossref | 100 | 0 | 100% |  |
| GBIF | 173 | 0 | 100% |  |
| INaturalist | 41 | 0 | 100% |  |
| IUCN | 95 | 29 | 69% | 81-83, 136-159, 162, 165 |
| NCBI | 104 | 0 | 100% |  |
| OBIS | 168 | 0 | 100% |  |
| WORMS | 104 | 0 | 100% |  |
| ZooBank | 73 | 0 | 100% |  |
| Total | 1020 | 29 | 97% |  |

### 2.6 Dependencies of *biodumpy*

The *biodumpy* package relies on several Python libraries, listed below: *Biopython* (v. 1.84): (Cock et al., 2009); *numpy* (v. 1.26.4): (Harris et al., 2020); *tqdm* (v. 4.66.4): (Casper da Costa-Luis et al., 2024); *requests* (v. 2.32.3): (Reitz, 2024); *lxml* (v. 5.2.2): (Behnel et al., 2024); *pandas* (v. 2.2.2) (The pandas development team, 2020); *beautifulsoup4* (v. 4.12.3): (Richardson, 2024); *pytest* (v. 8.3.3): (Krekel et al., 2004).

## 3 EXAMPLE APPLICATION

To demonstrate the basic usage of *biodumpy*, we provide the commented code of three comprehensive examples in *Supplementary Material 2*, which can be useful for introducing the potential of this new tool.

In the “Example 1”, we aim to work with several *biodumpy* modules and create a general pipeline to retrieve basic ecological information as occurrences and bibliography from a list of taxa. More specifically, we created a simple pipeline to store in a single file the taxonomic classification about a target species, its occurrences and an environmental variable (i.e., bathymetry) across a specified geographic area. Moreover, we retrieve the bibliographic information available in Zoobank. We divided this example into five steps: i) loading the necessary packages; ii) creating a taxa list and downloading information from GBIF, OBIS, COL, and ZooBank; iii) filtering the species-specific information and creating separate datasets containing taxonomy, distribution, and ZooBank bibliographic information; iv) saving each dataset into separate Excel sheets; v) creating two simple plots one to show the distribution of occurrences and another to depict the environmental variable.

In the “Example 2”, we focus on genetic modules, and we illustrate how to download FASTA files and perform an alignment with MAFFT (Katoh et al., 2019). To do that, we divided the process in five steps: i) loading the necessary packages; ii) creating a function to remove duplicated in shared records between BOLD and NCBI module outcomes; iii) downloading the FASTA files using BOLD and NCBI modules; iv) extracting only the sequences labeled as COI and merging the data into a single dataset; v) performing an alignment using MAFFT and save the result.

In the “Example 3”, we demonstrate how to use *biodumpy* within the R environment. This can be particularly useful for users not strictly familiarized with Python. To facilitate the integration of the package in R, we utilize the *reticulate* package (Ushey et al., 2024). We divide the example into three steps: i) install and load the reticulate package; ii) create a virtual environment in R; iii) use two *biodumpy* modules to download the data.

While the examples provided are useful for showing the potential of *biodumpy*, they do not fully capture the complexity or scope required for a comprehensive biological or ecological data management and analysis. Therefore, they should be considered as a simple guide to help the users in performing their own customized pipeline. A Python script and the downloaded data for each example are available at: Example link.

## 4. DISCUSSION

Recent advancements in computational power, coupled with the growing application of big data in various ecological domains, have significantly improved our capacity to analyse and understand the natural environment. Here, we introduced *biodumpy*, a novel and customizable Python package developed for the retrieval of biological data from multiple public databases. Unlike existing tools, which are often tailored to interact with a single data source, *biodumpy* includes several modules that enable users to download, manage, and integrate diverse types of biological information, primarily starting from a user-defined list of taxa. With the ability to retrieve different kinds of information simultaneously, *biodumpy* streamlines data collection and ensures compatibility across datasets, providing a robust foundation for ecological and biological analyses.

Overall, *biodumpy* represents a significant step toward enhancing the efficiency of data acquisition in ecological research, promoting interdisciplinary studies and enabling scientists to focus more on analysis and interpretation rather than on data download.

Future versions of the package will enhance flexibility for more biological data sources, integrate advanced data cleaning functions, and optimize performance for large-scale data processing.

## ACKNOWLEDGMENTS

This work has been partially funded and promoted by the Comunitat Autonoma de les Illes Balears through the Conselleria d’Educació i Universitats and by the European Union-Next Generation EU/PRTR-C17. I1 (SINCO2022/6717). Nevertheless, the views and opinions expressed are solely those of the author or authors, and do not necessarily reflect those of the Conselleria d’Educació i Universitats, the European Union or the European Commission.

Therefore, none of these organizations shall not be held liable.

This study has been partially funded by GOIB/Conselleria d’Educació i Universitats through the project “SINCO2022/18146” and co-funded by the European Union.

## AUTHOR CONTRIBUTIONS

CT and GDT conceived the ideas, designed methodology, create the documentation and test the software. CT wrote the original draft and led the writing of the manuscript. FA and RA designed methodology, tested the software, reviewed and edited the manuscript. CM reviewed and edited the manuscript. All authors contributed critically to the final version of manuscript and gave final approval for publication.

## DATA ACCESSIBILITY

OSF (Python scripts): https://osf.io/tn3au/?view_only=4ba55ec76e364afaa71f70778e0e69e5 GitHub: The code is available at: https://github.com/centrebalearbiodiversitat/biodumpy. To report a bug or issue, please contact the corresponding author or open an issue or pull request on the GitHub project.

