## Supplementary Material 1 for "*biodumpy*: A Comprehensive Biological Data Downloader"

### Equally shared first authorship

[1]Centre Balear de Biodiversitat, Universitat de les Illes Balears, Palma, Spain

* Corresponding author: Tommaso Cancellario
 -
CBB - University of the Balearic Islands (Spain)

**Supplementary Material 1**

**S1.** Examples of output for the BOLD module. (A) data in JSON format; (B) FASTA format; (C) summarized data using the parameter summary = True and (D) its field description.

**S2.** Examples of output for the COL module. (A) data retrieved using the parameter `check_syn` = *True*; (B) data retrieved setting the parameter `check_syn` = *False.*

**S3.** Examples of output for the Crossref module. (A) standard retrieved data; (B) summarized data and (C) its field description.

**S4.** Examples of output for the GBIF module. (A) data retrieved using the parameter `accepted_only` = *True*; (B) data retrieved using the parameter `accepted_only` = *False*; (C) occurrences downloaded within a specific polygon.

**S5.** Examples of output for the INaturalist module. (A) standard retrieved data.

**S6.** Examples of output for the IUCN module. (A) data retrieved using a list of regions; (B) data retrieved using the parameter `assess_details` = *True.*

**S7.** Examples of output for the NCBI module. (A) standard retrieved data; (B) data retrieved in FASTA format using the parameters `retype` = *"fasta"* `and output_format` = *fasta*; (C) summarized data using the parameter `summary` = *True* and (D) its field description.

**S8.** Examples of output for the OBIS module. (A) standard retrieved data; (B) occurrence data retrieved setting the `occ` = *True*, `geometry` = *'POLYGON((0.248 37.604, 6.300 37.604, 6.300 41.472, 0.248 41.472, 0.248 37.604))'*, and `areaid` = *33322.*

**S9.** Examples of output for the WORMS module. (A) standard retrieved data; (B) data retrieved coupled with the distribution information setting the parameter `distribution` = *True*.

**S10.** Examples of output for the ZooBank module. (A) data retrieved setting the parameter `dataset_size` = *"small"*; (B) data retrieved setting the parameter `dataset_size` = *"large"*; (C) data retrieved setting the parameter `dataset_size` = *"small"* and `info` = *True.*

**S11.** Commented Python script to create your own custom module.

**S1.** Examples of output for the BOLD module. (A) data in JSON format; (B) FASTA format; (C) summarized data using the parameter `summary` = *True* and (D) its field description.

**(A)** Example of data in JSON format: BOLD_A
**(B)** Example of data in FASTA format: BOLD_B
**(C)** Example of summarized data: BOLD_C
**(D)** Summarized data field description.

| Field name | Description |
|---|---|
| *record_id* | The unique identifier for the BOLD record. |
| *processid* | The process ID associated with the BOLD record. |
| *bin_uri* | The BIN (Barcode Index Number) URI. |
| *taxon* | The name of the lower taxonomic level. |
| *country* | The country where the collection event took place. |
| *province_state* | The province or state of the collection event. |
| *region* | The region of the collection event. |
| *lat* | The latitude of the collection event. |
| *lon* | The longitude of the collection event. |
| *markercode* | The marker code from the sequences data. |
| *genbank_accession* | The GenBank accession number from the sequences data. |

**S2.** Examples of output for the COL module. (A) data retrieved using the parameter `check_syn` = *True*; (B) data retrieved with the parameter `check_syn` = *False.*

**(A)** Example of data retrieved using the parameter `check_syn` = *True*: COL_A
**(B)** Example of data retrieved with the parameter `check_syn` = *False*: COL_B

**S3.** Examples of output for the Crossref module. (A) standard retrieved data; (B) summarized data and (C) its field description.

**(A)** Example of standard retrieved data: Crossref_A
**(B)** Example of summarized data: Crossref_B
**(C)** Summarized data field description.

| Field name | Description |
|---|---|
| *publisher* | The name of the publishing entity responsible for releasing the publication. |
| *container-title* | The title of the journal or book in which the research is published. |
| *DOI* | The Digital Object Identifier assigned to the publication. |
| *type* | The publication type, such as article, book chapter, report, etc. |
| *language* | The language in which the publication is written. |
| *URL* | A direct link to the publication. |
| *published* | The date when the research was published. |
| *title* | The title of the publication. |
| *author* | The names of the authors who contributed to the research along with their main academic information. |
| *abstract* | The publication abstract. |

**S4.** Examples of output for the GBIF module. (A) data retrieved using the parameter `accepted_only` = *True*; (B) data retrieved using the parameter `accepted_only` = *False*; (C) occurrences downloaded within a specific polygon.
`geometry` = *POLYGON((0.248 37.604, 6.300 37.604, 6.300 41.472, 0.248 41.472, 0.248 37.604))*

**(A)** Example of data retrieved using the parameter `accepted_only` = *True*: GBIF_A
**(B)** Example of data retrieved using the parameter `accepted_only` = *False*: GBIF_B
**(C)** Example of occurrence download: GBIF_C

**S5.** Examples of output for the INaturalist module. (A) standard retrieved data.

**(A)** Example of standard retrieved data: INaturalist_A

**S6.** Examples of output for the IUCN module. (A) data retrieved using a list of regions; (B) data retrieved using the parameter `assess_details` = *True*.

**(A)** Example of data retrieved using a list of regions: IUCN_A
**(B)** Example of data retrieved using the parameter `assess_details` = *True*: IUCN_B

**S7.** Examples of output for the NCBI module. (A) standard retrieved data; (B) data retrieved in FASTA format using the parameters `retype` = *"fasta"* and `output_format` = *fasta*; (C) summarized data using the parameter `summary` = *True* and (D) its field description.

**(A)** Example of standard retrieved data: NCBI_A

**(B)** Example of data retrieved in FASTA format: NCBI_B
**(C)** Example of summarized data: NCBI_C
**(D)** Summarized data field description.

| Field name | Description |
|---|---|
| *Id* | A numerical identifier (GI Number) that used to be assigned to each sequence version (e.g., "345678912"). |
| *Caption* | A unique identifier (accession number) assigned to a sequence when it is submitted to GenBank (e.g., "NM_001256789"). |
| *Title* | A short description or title of the sequence, often including information about the gene, organism, and type of sequence. |
| *Length* | The length of the sequence in base pairs (for nucleotide sequences) or amino acids (for protein sequences). |
| *query* | The original search term or query string used to retrieve this result. |

**S8.** Examples of output for the OBIS module. (A) standard retrieved data; (B) occurrence data retrieved setting the `occ` = *True*, `geometry` = *'POLYGON((0.248 37.604, 6.300 37.604, 6.300 41.472, 0.248 41.472, 0.248 37.604))'*, and `areaid` = *33322*.

**(A)** Example of standard retrieved data: OBIS_A
**(B)** Example of occurrence download: OBIS_B

**S9.** Examples of output for the WORMS module. (A) standard retrieved data; (B) data retrieved coupled with the distribution information setting the parameter `distribution` = *True*.

**(A)** Example of standard retrieved data: WORMS_A
**(B)** Example of data retrieved with distribution information: WORMS_B

**S10.** Examples of output for the ZooBank module. (A) data retrieved setting the parameter `dataset_size` = *"small"*; (B) data retrieved setting the parameter `dataset_size` = *"large"*; (C) data retrieved setting the parameter `dataset_size` = *"small"* and `info` = *True.*

**(A)** Example of data retrieved setting the parameter `dataset_size` = *"small"*: ZooBank_A
**(B)** Example of data retrieved setting the parameter `dataset_size` = *"large"*: ZooBank_B
**(C)** Example of data retrieved setting the parameter `dataset_size` = *"small"* and `info` = *True:* ZooBank_C

**S11.** Commented Python script to create your own custom module: Python_script
