## Supplementary Material 2 for "*biodumpy*: A Comprehensive Biological Data Downloader"

### Equally shared first authorship

[1]Centre Balear de Biodiversitat, Universitat de les Illes Balears, Palma, Spain

* Corresponding author: Tommaso Cancellario
 -
CBB - University of the Balearic Islands (Spain)

**Supplementary Material 2**

**Example 1 – Work with taxonomic, distributional, and bibliographic data**

In this example, we aimed to create a simple pipeline to store in a single file the systematics taxonomy classification about a target species, its occurrences and an environmental variable (i.e., bathymetry) across a specified geographic area, and the bibliographic information available in ZooBank for the given taxon. To achieve this, we divided the example into five steps: i) loading the necessary packages; ii) creating a taxa list and downloading information from GBIF, OBIS, COL, and ZooBank; iii) filtering the species-specific information and creating separate datasets containing taxonomy, distribution, and ZooBank bibliographic information; iv) saving each dataset into separate Excel sheets; v) creating two simple plots one to show the distribution of occurrences and another for the environmental variable.

Link: Example_1
Data accessed: April 23, 2025.

Firstly, we import the required package for conducting the analysis. If packages are not already installed in the Python virtual environment, they can be installed using the following command: `pip install package_name==version_number`.

```
# 1) Import packages and modules

import json
import statistics
import seaborn as sns
import matplotlib.pyplot as plt
import xlsxwriter

from biodumpy import Biodumpy
from biodumpy.inputs import GBIF, OBIS, COL, ZooBank
```

After imported the packages, we define a list of taxa to start the download. In this case, we used a single taxon (e.g., *Pinna nobilis*) to expedite the analysis. However, users can include a list of taxa as desired. We utilized the GBIF, OBIS, COL, and ZooBank modules to retrieve various types of information, such as taxon occurrences, taxonomy, and bibliography. To limit the occurrence download, we defined a polygon around the Balearic Islands.

```
# 2) Create a taxa list and download data

taxa = ['Pinna nobilis']
polygon = ‘POLYGON((1.1169 38.56042,4.41447 38.56042,4.41447 40.19769,1.116
9 40.19769,1.1169 38.56042))’

bdp = Biodumpy([
    GBIF(geometry=polygon, occ=True, accepted_only=True),
    OBIS(geometry=polygon, occ=True),
    COL(),
    ZooBank(dataset_size=‘small’, info=True)
])
bdp.start(taxa)
```

After downloading the data, we can filter each JSON file and create specific objects for each data type.

```

# Extract taxonomy of Pinna nobilis provided by COL

f = open('./COL/Pinna nobilis.json', 'r')
data_col = json.load(f)[0]
f.close()


classification = data_col[‘classification’]

taxonomy = []
for item in classification:
    taxonomy.append({
        'id': item['id'],
        'name': item['name'],
        'rank': item['rank']
    })

# Extract DOI
f = open('./ZooBank/Pinna nobilis.json', 'r')
data_zoo = json.load(f)
f.close()
doi = [item['info'][0]['IdentifierURL'] for item in data_zoo]


# 3) Prepare distributional data
f = open('./GBIF/Pinna nobilis.json', 'r')
data_gbif = json.load(f)
f.close()

f = open('./OBIS/Pinna nobilis.json', 'r')
data_obis = json.load(f)
f.close()
```

```
# Filter GBIF and OBIS data and merge into a single dataset
data_gbif_filtered = [
    {
        'scientificName': item.get('acceptedScientificName', None),
        'decimalLatitude': item.get('decimalLatitude', None),
        'decimalLongitude': item.get('decimalLongitude', None),
        'depth': item.get('depth', None),
        'source': 'GBIF'
    } for item in data_gbif
]

data_obis_filtered = [
    {
        'scientificName': item.get('scientificName'),
        'decimalLatitude': item.get('decimalLatitude'),
        'decimalLongitude': item.get('decimalLongitude'),
        'depth': item.get('bathymetry'),
        'source': 'OBIS'
    } for item in data_obis
]

data = data_gbif_filtered + data_obis_filtered

# Create an object with depth information
depth = [item['depth'] for item in data if item['depth']]
mean_depth = round(statistics.mean(depth), ndigits=2)
median_depth = round(statistics.median(depth), ndigits=2)
```

We now create a tabular file (e.g., .xlsx) containing the downloaded information, organized into separate sheets.

```
# 4) Save files into an .xlmx

workbook = xlsxwriter.Workbook('./biodumpy_ex_1.xlsx')
worksheet = workbook.add_worksheet('taxonomy')
worksheet. write_row('A1', ['Rank', 'Name'])

# Write taxonomy data
for i, item in enumerate(taxonomy, start=2):
    Worksheet.write_row(f'A{i}', [item['rank'], item['name']])

worksheet = workbook.add_worksheet('occurrences')

Worksheet.write_row('A1', ['scientificName', 'decimalLatitude',  'decimalLo
ngitude', 'depth', 'source'])

# Write occurrence data
for i, item in enumerate(data, start=2):
      worksheet.write_row(f'A{i}', [item['scientificName'], item['decimalLa
titude'], item['decimalLatitude'], item['depth'], item['source']])

worksheet = workbook.add_worksheet('bibliography')
worksheet.write('A1', 'doi')

# Write bibliographic data
for i, item in enumerate(doi, start=2):
    worksheet.write(f'A{i}', item)

# Save the workbook
```

```
workbook.close()
```

Finally, we produce two simple plots. The first is a density plot illustrating the depth data's distribution, along with its associated mean and median (Fig. 1). The second is a joint plot showing the relationship between latitude and longitude variables (Fig. 2).

```
# 5 Create density plot of depth and another joint plot for longitude and latitude data

sns.kdeplot(depth, color='orange', fill=True, alpha=.3, linewidth=0)
plt.axvline(x=mean_depth, color='blue', linestyle='--', linewidth=1, label='Mean')
plt.axvline(x=median_depth, color='purple', linestyle='--', linewidth=1, label='Median')
plt.legend()
plt.show()

lon = [item['decimalLongitude'] for item in data]
lat = [item['decimalLatitude'] for item in data]
sns.jointplot(x=lon, y=lat, kind='scatter', color='orange', edgecolor=‘black’)
plt.show()
```

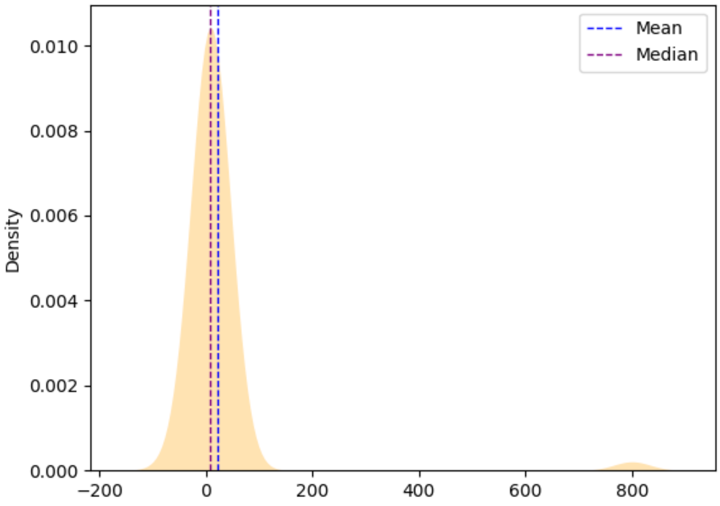


**Fig. 1.** The figure shows a density plot representing the distribution of depth data. The x-axis indicates the depth values, while the y-axis represents the corresponding density. A vertical dashed blue line marks the mean depth, whereas a purple line indicates the median depth. The shaded area under the curve highlights the overall density distribution.

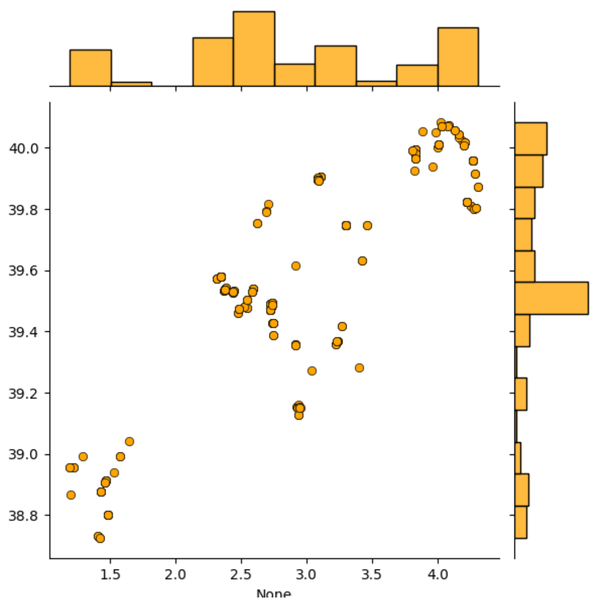


**Fig. 2.** The figure presents a joint plot illustrating the relationship between latitude and longitude variables. The central scatter plot depicts individual data points, with longitude values on the x-axis and latitude values on the y-axis, indicating the spatial distribution of occurrences. Marginal histograms are displayed along the axes, showing the frequency distributions of longitude and latitude independently.

**Example 2 – Genetics**

In this example, we focus on genetic modules, and we illustrate how to download FASTA files and perform an alignment with MAFFT. We divided the process in five main steps: i) loading the necessary packages; ii) creating a function to remove duplicate records; iii) downloading the FASTA files from BOLD and NCBI modules; iv) extracting only the sequences labeled as COI and merging the data into a single dataset; v) performing an alignment using MAFFT.

Link: [Example_2]
Data accessed: April 23, 2025.

First, we load the packages, and the modules needed to perform the analysis.

```
# 1) Import packages
import subprocess

from biodumpy import Biodumpy
from biodumpy.inputs import BOLD, NCBI
from biodumpy.utils import read_fasta, save_fasta
```

Next, we create a Python function to remove possible duplicate records shared between the two datasets. This function has two input parameters: `dicts_list`, which specifies the name of the list containing the sequences, and `key`, which indicates the name of the field containing the values to check for duplicates. In this example, we use the NCBI accession number as the value to check.

```
# 2) Create ‘remove_duplicates’ function
def remove_duplicates(dicts_list: list, key: str):
    seen = set()
    unique_dicts = []

    for d in dicts_list:
        value = d.get(key)
        if value not in seen:
            unique_dicts.append(d)
            seen.add(value)

    return unique_dicts
```

Now, we download the sequences in FASTA format for the Odonata species *Anax imperator* using the *biodumpy* modules BOLD and NCBI.

```
# 3) Create a taxa list
taxa = [‘Anax imperator’]

# Set the modules BOLD and NCBI and start the download
bdp = Biodumpy([
      BOLD(bulk=True, output_format='fasta'),
      NCBI(bulk=True, rettype='fasta', output_format='fasta',
      query_type='[Organism]')
])

bdp.start(taxa,output_path='./downloads/{date}/{module}/{name}')
```

Once we download the files, we can open the two FASTA files using the function `read_fasta`, specifying the file paths of the downloaded data as parameters. Next, we extract only the sequences labeled with COI and combine the data obtained from BOLD and NCBI modules into a single dataset removing duplicated records.

```
# 4) Extract only COI Marker
seq_bold = read_fasta('./downloads/{date}/BOLD/bold.fasta')
seq_ncbi = read_fasta('./downloads/{date}/NCBI/ncbi.fasta')

# Filtered only sequences labeled as COI
filtered_bold =
[item for item in seq_bold if 'COI-5P' in item['id']]
filtered_ncbi =
[item for item in seq_ncbi if 'cytochrome oxidase subunit 1' in item['id']]

# Remove shared records between bold and ncbi
for item in filtered_bold:
    item['ncbi_accession'] = item['id'].split('|')[-1]      if len(item['id
'].split('|')) == 4 else None

for item in filtered_ncbi:
    item['ncbi_accession'] = item['id'].split('.')[0]

# Join the filtered data
```

```
sequences = remove_duplicates( filtered_ncbi + filtered_bold, 'ncbi_accessi
on')

save_fasta('Saving Path', sequences)
```

Finally, we perform sequence alignment using MAFFT and save the resulting output. In this step, we create a command to run MAFFT, setting the alignment strategy parameter to *auto* (for more information about MAFFT parameter options, please visit the official webpage: https://mafft.cbrc.jp/alignment/software/).

```
# 5) Alignment
input_file = 'Path of the input.fasta'
output_file = 'Path of the output.fasta'

# Construct the MAFFT command
mafft_command = ['mafft', '--auto', input_file]

# Run MAFFT
with open(output_file, 'w') as output:
    subprocess.run(mafft_command, stdout=output)
```

**Example 3 – *biodumpy* in R**

In the following example, we describe the procedure for integrating *biodumpy* into the R environment. This example is run within the R environment. Therefore, the code syntax will be slightly different compared to the examples above. We divide the following code into three main steps: i) install and load the reticulate package; ii) create a virtual environment in R; iii) use a couple of *biodumpy* modules to download the data.

Link: Example_3
Data accessed: April 23, 2025.

As the first step we install and load the pack *reticulate*.

```
# 1) Install and load the reticulate pack
install.packages('reticulate', dependencies=TRUE)
library(reticulate)
```

Once the package is installed and loaded, users can set up a specific virtual environment to perform the analysis. While this step may be unfamiliar to many R users, it is important to understand its purpose. Similar to R, Python has a vast ecosystem of external packages that extend its core functionality. However, Python allows only one version of a package to be installed at a time, which can create conflicts when working on multiple projects requiring different package versions or dependencies. To address this issue, Python enables the creation of virtual environments, which allow users to manage packages and their dependencies independently for each project. By default, *reticulate* uses an isolated Python virtual environment named ‘r-reticulate’.

```
# 2) Create a new environment
virtualenv_create('example_biodumpy')

# Set a specific virtualenv and install biodumpy
use_virtualenv('example_biodumpy')
virtualenv_install('example_biodumpy', 'biodumpy')
```

Once we set the virtual environment containing the *biodumpy* package it is possible to start to work with it. Similarly to the previous examples, we download occurrences and sequences from GBIF and BOLD respectively.

```
# 3) Import biodumpy and its modules
biodumpy <- import('biodumpy')
inputs <- import('biodumpy.inputs')

# Define the taxa vector
taxa <- list('Anax imperator')

# Define a polygon to limit the download from GBIF
polygon = 'POLYGON((1.1169 38.56042,4.41447 38.56042,4.41447 40.19769,1.116
9 40.19769,1.1169 38.56042))'

# Set the BOLD and GBIF modules
bold <- inputs$BOLD(bulk=FALSE, fasta=TRUE, output_format='fasta')
gbif <- inputs$GBIF(bulk=FALSE, occ = TRUE, geometry = polygon)

# Start the download
bdp <- biodumpy$Biodumpy(list(bold, gbif))
bdp$start(taxa, output_path = './{date}/{module}/{name}')
```
